# Meta analysis of variant predictions in congenital adrenal hyperplasia caused by mutations in CYP21A2

**DOI:** 10.1101/2021.12.21.473700

**Authors:** Mayara J. Prado, Rodrigo Ligabue-Braun, Arnaldo Zaha, Maria Lucia Rosa Rossetti, Amit V. Pandey

## Abstract

**Context:** CYP21A2 deficiency represents 95% of congenital adrenal hyperplasia cases (CAH), a group of genetic disorders that affect steroid biosynthesis. The genetic and functional analysis provides critical tools to elucidate complex CAH cases. One of the most accessible tools to infer the pathogenicity of new variants is *in silico* prediction.

**Objective:** Analyze the performance of *in silico* prediction tools to categorize missense single nucleotide variants (SNVs) of the CYP21A2.

**Methods:** SNVs of the CYP21A2 characterized *in vitro* by functional assays were selected to assess the performance of online single and meta predictors. SNVs were tested separately or in combination with the related phenotype (severe or mild CAH form). In total, 103 SNVs of the CYP21A2 (90 pathogenic and 13 neutral) were used to test the performance of 13 single-predictors and four meta-predictors.

**Results:** SNVs associated with the severe phenotypes were well categorized by all tools, with an accuracy between 0.69 (PredictSNP2) and 0.97 (CADD), and Matthews’ correlation coefficient (MCC) between 0.49 (PoredicSNP2) and 0.90 (CADD). However, SNVs related to the mild phenotype had more variation, with the accuracy between 0.47 (S3Ds&GO and MAPP) and 0.88 (CADD), and MCC between 0.18 (MAPP) and 0.71 (CADD).

**Conclusion:** From our analysis, we identified four predictors of CYP21A2 pathogenicity with good performance. These results can be used for future analysis to infer the impact of uncharacterized SNVs’ in CYP21A2.

## Introduction

One of the most common autosomal recessive genetic disorders is the impairment of the steroid 21-hydroxylase (CYP21A2). This CYP21A2 deficiency represents about 95% of cases in the congenital adrenal hyperplasia (CAH), a group of enzymatic disorders that affect cortisol biosynthesis. The CYP21A2 enzyme is a member of the cytochrome P450 superfamily (CYPs) and catalyzes the conversion of 17α-hydroxyprogesterone (17OHP) into 11-deoxycortisol and progesterone into 11-deoxycorticosterone. Other enzymes subsequently convert these steroids into cortisol and aldosterone, respectively ^1^.

Clinically, the CYP21A2 deficiency in humans has a wide spectrum of phenotypes, from severe to mild or asymptomatic ^2,3^. The classical severe CAH has salt-wasting (SW) and simple-virilizing (SV) forms. The classical SW form has no enzyme activity and is related to severe virilization and electrolyte dysregulation. In contrast, the classical SV form has enough residual enzyme activity to prevent adrenal crisis ^2^. The mild CAH is the non-classical (NC) form of CAH and has a partial CYP21A2 activity associated with hyperandrogenism and mild late-onset CAH ^3^. Furthermore, there is a relatively good genotype-phenotype correlation for CYP21A2 deficiency, which allows the categorization of variants according to the residual enzyme activity (obtained from *in vitro* studies) and their expected phenotype ^3^. The classical CAH has less than 10% of wild-type (WT) enzyme activity in 95% of the cases, while the NC form has an activity between 10-78% of the WT in 90% of the cases. ^4^.

The *CYP21A2* gene is a tandemly arranged module (RCCX: *RP-C4-CYP21*-*TNX*) and shows 96-98% of sequence identity with its pseudogene, *CYP21A1P* ^5^. These features make the *CYP21A2* gene analysis a complex endeavor, with many different types of mutations - from single nucleotide variants (SNVs) to genetic rearrangements - and further complicated by the fact that most carriers have compound heterozygous mutations ^3^. However, only ten mutations described in the general population are sampled by CYP21A2 deficiency screening programs. The whole gene sequence analysis by Sanger sequencing is an alternative method for exceptional cases due to the cost and time-consuming nature of such studies ^6,7^.

So far, with the whole *CYP21A2* gene sequencing, genetic studies have reported more than 1,300 variants for the *CYP21A2* gene. Out of the 230 variants reported as affecting human health, 153 are missense variants ^4^. The advancement of next-generation sequencing (NGS) to analyze a large number of genes has facilitated the detection of rare single nucleotide variants (SNVs) and single nucleotide polymorphisms (SNPs). A few years ago, this technology was not applied to screen the *CYP21A2* gene defects due to its high sequence identity with *CYP21A2P* which hampers the proper analysis of this genomic region ^5^. However, recently some groups have found alternative ways to perform NGS for the *CYP21A2* gene through a combination with other methods, such as Multiplex ligation-dependent probe amplification ^8,9^. These genetic analysis strategies of the *CYP21A2* gene with the NGS technology represent a promising tool for the future, opening the window to identify new variants while improving the diagnosis of CYP21A2 deficiency, and establishing a more reliable estimate of mutation frequencies.

The gold standard for the characterization of new CYP21A2 variants is *in vitro* functional assay. However, this approach takes too much time, and it is not viable for all-new variants detected. One of the most accessible tools to predict the pathogenicity of variants is the *in-silico* analysis, which usually has free access, a friendly interface, and provides quick results. Many online predictors are available that have different features and approaches, from single characteristic analysis to meta-predictors with different compositions and algorithms. Some studies have shown the general performances of these tools against a whole database with few predictors ^10,11^. However, studies with variants on protein-specific analysis showed that general analysis results cannot be extrapolated for all proteins as each protein has unique characteristics, which is a key limitation of predictor programs ^12–14^. Therefore, it is essential to be careful when choosing the prediction tools and to consider their variable accuracies for each gene ^11^.

Here we have done a meta-analysis for the performance of online predictor tools to classify missense SNVs of the CYP21A2. Missense SNV is the most common group of variants in the human genome, one at every kilobase. In the *CYP21A2* gene, **this type of SNV** represents about 60% of the CYP21A2 variants in The Human Gene Mutation Database (HGMD, RRID:SCR_001888)^6^ and 65% of those affecting human health ^4^. Additionally, missense SNV is one of the hardest variant types for interpretation ^4,15,16^. In total, we analyzed 17 predictors with multiple algorithms, approaches and datasets. Thirteen of these were based on single features: CADD (RRID:SCR_018393)^17^, ConSurf (RRID:SCR_002320) ^18^, DANN ^19^, FATHMM ^20^, MAPP (RRID:SCR_014375) ^21^, MutPred2 (RRID:SCR_010778) ^22^, PANTHER-PSEP (RRID: SCR_005145) ^23^, PhD-SNP^g^ (RRID: SCR_010782) ^24^, PolyPhen-2 (RRID:SCR_013189) ^25^, PROVEN (RRID: SCR_002182) ^26^, SIFT (RRID:SCR_012813) ^27^, SNAP2 (RRID:SCR_002127) ^28^, and SNPs&GO (RRID:SCR_005788)^29^. Four meta-predictors: PredictSNP ^30^, PredictSNP2 ^31^, Meta-SNP ^32^, and S3Ds&GO ^33^) (Figure 1). We excluded nonsense and frameshift variants from our analysis since they have specific settings in many predictors and a high agreement ratio between tools.

**Figure 1.**
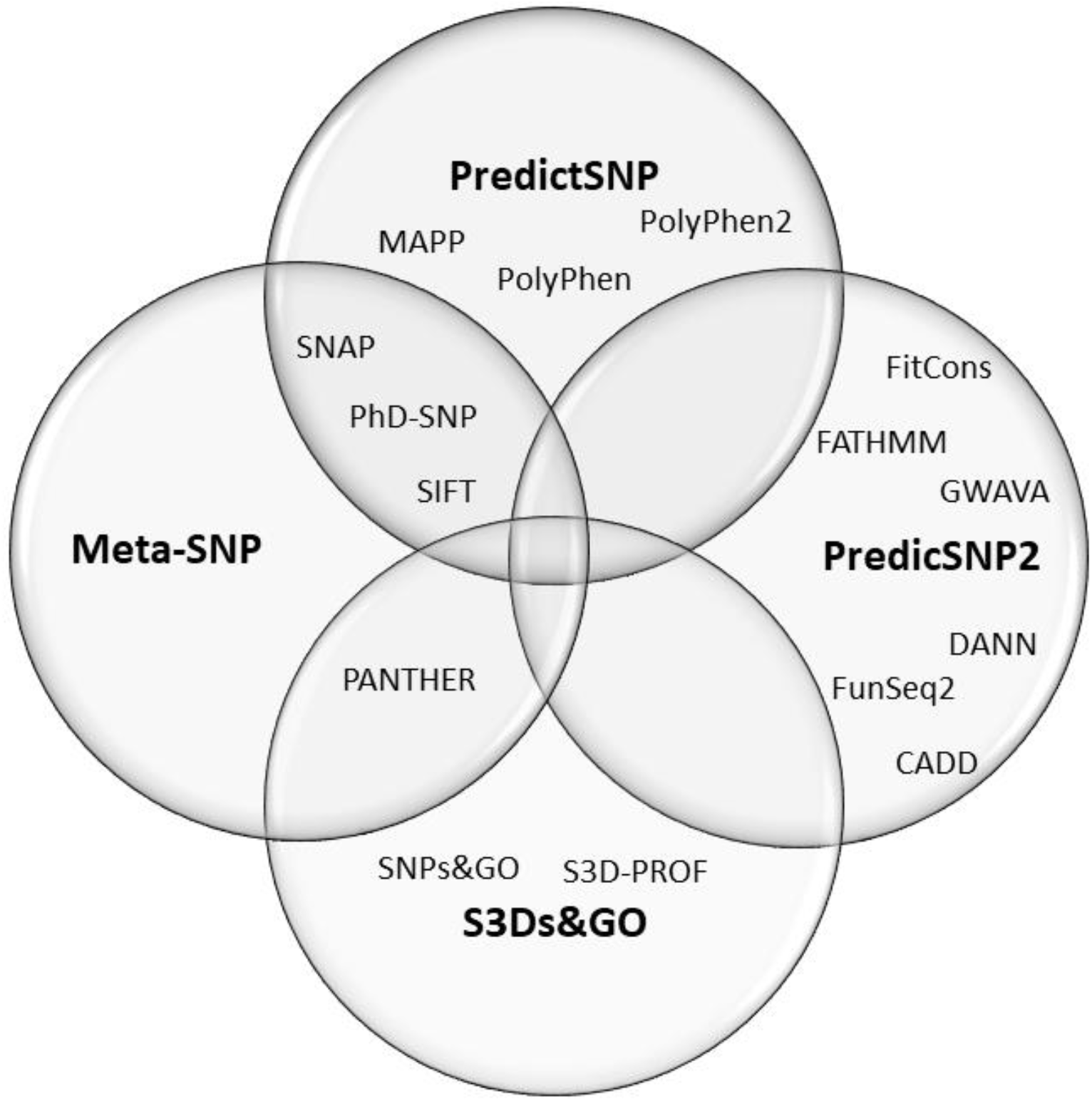
Composition of the four meta-predictors studied. The PredictSNP algorithm comprises outputs of six single-predictors, the PredictSNP2 of six, the Meta-SNP of 4, and the S3Ds&GO of three predictors.

**Figure 2.**
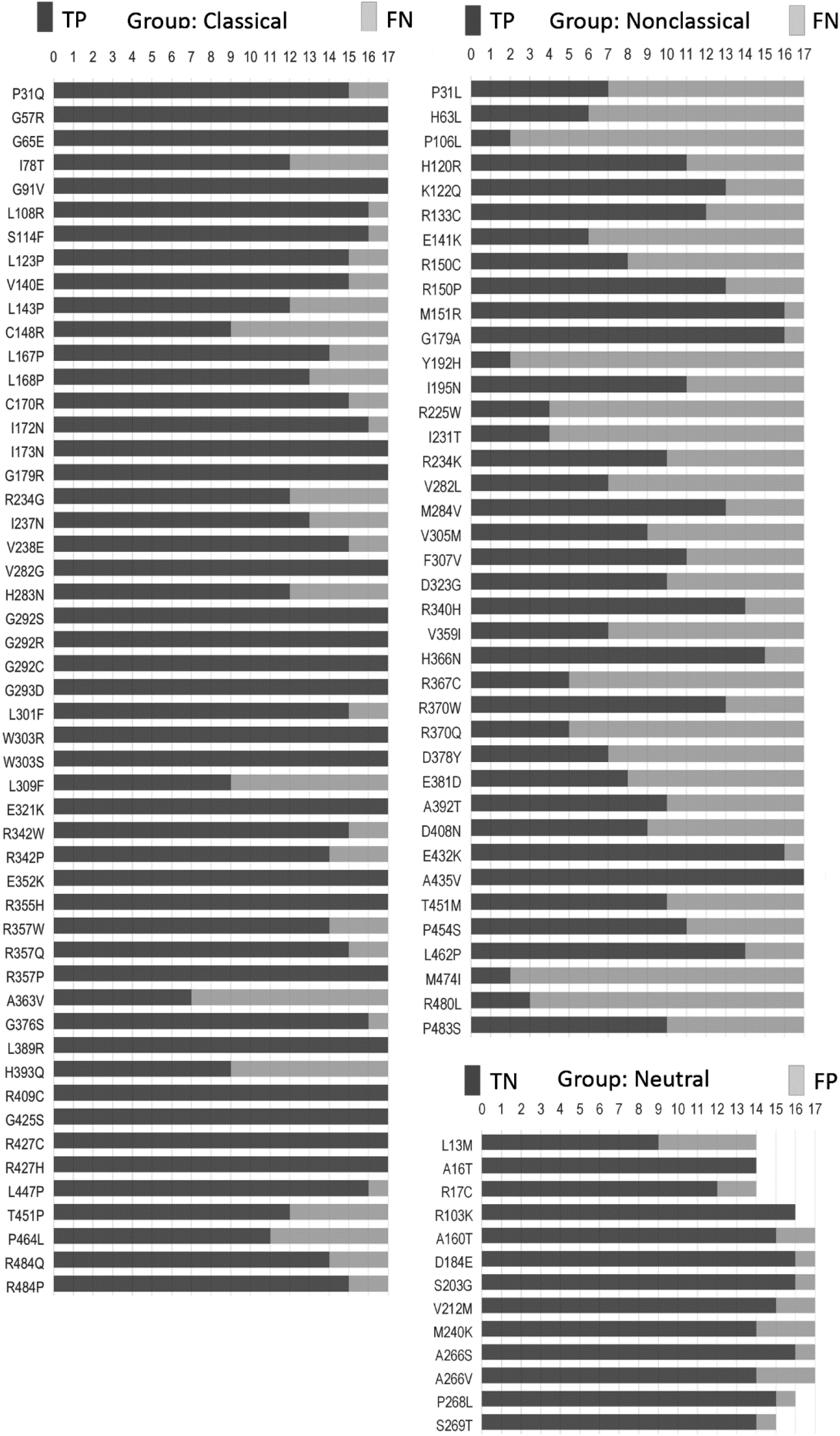
Frequency of hit and miss obtained for each SNV by mutation group. Each SNV (vertical list) was analyzed by seventeen predictors (horizontal measurement) performed with the default setting for missense mutation. Please, refer to Table S4-S6 for details. TP, true positive; TN, true negative; FN, false negative; FP, false positive.

## Materials and Methods

### SNVs selection and categorization

To select CYP21A2 missense single nucleotide variants (SNVs) with clinical significance, we used the list of variants reported to affect human health, as reviewed by *Simonetti et al* ^4^. Complementarily, we searched for SNVs reported by dbSNP, Ensembl, and GeneCards applying the following filter when present: “missense”, clinical significance “pathogenetic” or “benign”, “without conflicting interpretation”, and “human or homo sapiens”. We excluded nonsense and frameshift mutations. In addition, we performed a cross-check of the databases with original articles or reviews to remove variants without the enzyme activity data available.

To standardize the effect of the SNVs selected, we categorized them into three groups according to the CYP21A2 activity for at least one of the steroid substrates: classical (CL), nonclassical (NC), and neutral. The CL group has SNVs with the CYP21A2 activity level < 10 % relative to WT, the NC group has SNVs with the activity level between > 10% and < 78 % relative to WT, and the neutral group has SNV with the enzyme activity > 78 % relative to WT. The CAH group is composed of all SNVs from the CL and NC. The mean and standard deviation (SD) of enzyme activity were calculated for each steroid and mutation group.

### Selection of predictor tools

To choose predictors with different features, we reviewed the literature for software applied to *in silico* analysis of SNP or SNVs. Predictors used in more than two studies by different research groups or significant performance in a large study were selected. In addition, we filtered for tools with free access and online availability, thus not requiring local powerful computational resources. Characteristics of each predictor chosen are shown in Table S1-S2.

### Data treatment

The default setting for missense mutation was used on all predictors. However, when there was no set instruction for that purpose highlighted for the program, we followed the developers’ recommendation from the tutorial or original paper. In addition, three scores were extracted indirectly from meta-predictors: MAPP (v.28.6.2005) and SIFT (v.4.0.4) scores were obtained from PredictSNP, and DANN (v.1.2) score from PredictSNP2. For statistical purposes, we standardized two variables for the outputs of all the predictors: “**Damage**” for SNVs with the potential to affect CYP21A2 and “**neutral**” for SNVs with no or very low potential to affect the enzyme. The following outputs were standardized as “damage”: score > 0.5 to Meta-SNP, SNP&GO, S3D&GO, MutPred2, FATHMM-MKL (weighted) and PhD-SNP^g^; “deleterious” message to PredictSNP, PredictSNP2, MAPP, and SIFT; score > 0.9 to DANN; score > -2.5 to PROVEN; score > 10 to CADD (GRCh38-v1.5-6); score > 0.45 PolyPhen-2 (HumVar); score < 0 to ConSurf; score > 0 to SNAP2; and score > 450 millions of years to PANTHER. Otherwise, we classified the outputs as “neutral”.

### Analytical parameters

We analyzed the performance of each predictor in two ways. First, to assess the performance to discriminate the effects of *CYP21A2* SNVs, we compared SNVs of the CAH group with the neutral group. Second, to get the number of hits and misses per group, we analyzed CL and NC groups separated against the neural group. We used Microsoft Excel for the data organization and, together with IBM SPSS Statistic software v.2.1, we performed the statistical analysis.

### Statistical methods

We considered true positive (TP) result for correct “damage” prediction, true negative (TN) for correct “neutral” prediction, false positive (FP) for incorrect “neutral” prediction, and false-negative (FN) for incorrect “damage” prediction. We calculated the positive predictive values (PPV) to access the ratio of TP results for all positive results (Eqn. 1), and negative predictive values (NPV)NPV to the ratio of TN for all negative results (Eqn. 2).

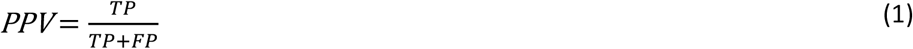

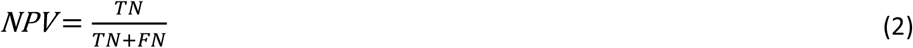

The proportion of correct SNVs identified as harmful was assessed with the sensibility (Se) equation (Eqn. 3), while the correct neutral identification was assessed with the specificity (Sp) (Eqn. 4). Besides that, we obtained the accuracy (Ac) by the ratio of true results (TP and TN) (Eqn. 5). The accuracy (Ac) was classified as excellent (0.9 < Ac < 1.0), good (0.8 < Ac < 0.9), fair (0.7 < Ac < 0.8), and not good (0.6 < Ac < 0.7).

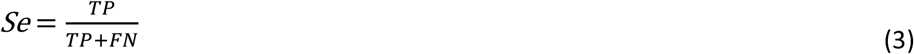

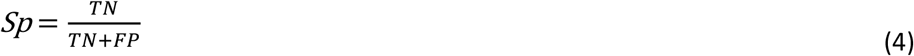

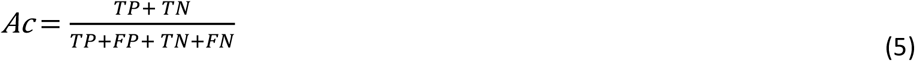

Finally, we applied the MCC to measure the two-class quality (harmful and neutral). This method is suitable for imbalanced data and has been used to evaluate *in silico* prediction approaches. MCC score ranges from 1 (perfect prediction) to -1 (total disagreement between the results predicted and observed), with 0 being no better than random prediction (Eqn. 6) (Chicco et al., 2021).

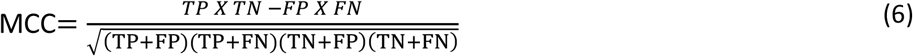

## Results

### Data of the selected SNVs

From variants in the *CYP21A2* gene reported in the literature and databases, we selected missense SNVs with clinical significance (criteria described in section 4.1). We obtained 96 valid SNVs out of 299 missense variants in the list [4], 85 out of 614 missense variants in dbSNP, 66 out of 459 missense variants in Ensembl, 45 out of 71 missense variants in GeneCards, 47 out of 83 missense variants in ClinVar, 26 in OMIM, and 81 in UniProt databases.

By removing SNVs that were duplicated and with no functional characterization, we obtained 103 SNVs, 51 classified as classical, 39 as non-classical, and 13 as neutral. The SNVs selected with the respective enzyme activity are described in table S3. All studies presented the CYP21A2 activity measured by the hydroxylation of 17OHP, while 86 also measured progesterone hydroxylation. Mutations of the CL group have the mean enzyme activity for the 17OHP hydroxylation of 1.5 ± 2 (SD)% and the progesterone hydroxylation of 1.3 ± 1.6%. While mutations of the NC group have 17OHP hydroxylation activity of 42.9 ± 23% and progesterone hydroxylation activity of 37.4 ± 21%. Finally, mutations in the neutral group have 17OHP hydroxylation activity of 100.17 ± 11% and progesterone hydroxylation activity of 94.1 ± 8.25%.

### General analysis of mutation groups

We obtained 22 SNVs in the *CYP21A2* gene with the correct prediction for all tested predictors, although the exact number was incorrectly predicted for at least half of them (Table S3). There was no SNV wrongly predicted by all predictors. We compared the hit and miss by the 17 predictors for all SNV affecting CYP21A2 activity, the CAH group, and for all the non-pathogenic SNV of the neutral group. We showed that 22% (22 of 90) SNVs from the CAH group obtained the correct score by all predictors, while 23% (21 of 90) were wrongly predicted by at least nine tools. The neutral group got 2 of its 13 SNVs (15.4%) rightly mis predicted by all and one by nine predictors. Moreover, we divided the SNV of the CAH group into the CL and NC groups. We got 37% (out of 51 SNVs) from the CL and 2.6% (of 39 SNVs) from the NC groups SNVs correctly predicted by all tools. While 5.9% and 46% from CL and NC groups, respectively, were wrongly predicted by nine tools.

### Performance of predictors to identify SNVs detrimental to CYP21A2 activity

We analyzed the performance of 17 predictors to identify the 90 SNVs that affect CYP21A2 functionality against the 13 SNVs with neutral effect (Table 1). All the 17 predictors obtained a good PPV rate (> 0.90). However, only CADD (0.73) and DANN (0.56) showed negative predictive values (NPVs) higher than 0.5. PANTHER-PSEP has no result for the NPV, as it could not identify benign variants.

**Table 1.** Performance of 17 programs to predict the effect SNVs in the *CYP21A2*. We performed the analysis with 103 functionally characterized variants, 90 damaging the protein functionality, and 13 neutral—Color scores from blue (good result) to yellow (not good).

| Predictors | TP | FN | TN | FP | PPV | NPV | Se | Sp | Ac | MCC |
| --- | --- | --- | --- | --- | --- | --- | --- | --- | --- | --- |
| PredictSNP<sup>a</sup> | 58 | 32 | 13 | 0 | 1.00 | 0.29 | 0.64 | 1.00 | 0.69 | 0.43 |
| PredictSNP2<sup>a</sup> | 44 | 46 | 13 | 0 | 1.00 | 0.22 | 0.49 | 1.00 | 0.55 | 0.33 |
| Meta-SNP<sup>a</sup> | 60 | 30 | 13 | 0 | 1.00 | 0.30 | 0.67 | 1.00 | 0.71 | 0.45 |
| S3Ds&GO<sup>a</sup> | 58 | 32 | 8 | 0 | 1.00 | 0.20 | 0.64 | 1.00 | 0.67 | 0.36 |
| CADD | 86 | 4 | 11 | 2 | 0.98 | 0.73 | 0.96 | 0.85 | 0.94 | 0.75 |
| ConSurf | 79 | 11 | 9 | 1 | 0.99 | 0.45 | 0.88 | 0.90 | 0.88 | 0.58 |
| DANN | 83 | 7 | 9 | 4 | 0.95 | 0.56 | 0.92 | 0.69 | 0.89 | 0.56 |
| FATHMM | 62 | 28 | 13 | 0 | 1.00 | 0.32 | 0.69 | 1.00 | 0.73 | 0.47 |
| MAPP | 54 | 36 | 10 | 2 | 0.96 | 0.22 | 0.60 | 0.83 | 0.63 | 0.28 |
| MutPred2 | 62 | 28 | 13 | 0 | 1.00 | 0.32 | 0.69 | 1.00 | 0.73 | 0.47 |
| PANTHER-PSEP | 90 | 0 | 0 | 9 | 0.91 | nr | 1.00 | 0.00 | 0.91 | nr |
| PhD-SNPg | 62 | 28 | 12 | 1 | 0.98 | 0.30 | 0.69 | 0.92 | 0.72 | 0.42 |
| PolyPhen2-HumVar | 79 | 11 | 12 | 1 | 0.99 | 0.52 | 0.88 | 0.92 | 0.88 | 0.64 |
| PROVEAN | 66 | 24 | 13 | 0 | 1.00 | 0.35 | 0.73 | 1.00 | 0.77 | 0.51 |
| SIFT | 70 | 20 | 11 | 2 | 0.97 | 0.35 | 0.78 | 0.85 | 0.79 | 0.45 |
| SNP2 | 59 | 31 | 13 | 0 | 1.00 | 0.30 | 0.66 | 1.00 | 0.70 | 0.44 |
| SNPs&GO | 54 | 36 | 13 | 0 | 1.00 | 0.27 | 0.60 | 1.00 | 0.65 | 0.40 |

<sup>a</sup> meta-predictor. nr, no result; PPV, positive predictive value; NPV, negative predictive value; Se, sensibility; Sp, specificity; Ac, accuracy; MCC, Matthews' correlation coefficient test.

**Table 2.** Performance of 17 programs for the specific CYP21A2 groups. We predict the effect of SNVs in the CYP21A2 dividing them by the two levels of protein damage: the severe (classical mutation, CL group) and mild (non-classical mutation, NC group). We performed the analysis with 103 SNVs of known effect, 51 being CL, 39 NC, and 13 neutral. Color score from blue (good result) to yellow (not good).

| Specifics groups | PPV |  | NPV |  | Se |  | Sp |  | Ac |  | MCC |  |
| --- | --- | --- | --- | --- | --- | --- | --- | --- | --- | --- | --- | --- |
|  | CI | NC | CI | NC | CI | NC | CI | NC | CI | NC | CI | NC |
| PredictSNP<sup>a</sup> | 1.00 | 1.00 | 0.65 | 0.34 | 0.86 | 0.36 | 1.00 | 1.00 | 0.89 | 0.52 | 0.75 | 0.35 |
| PredictSNP2<sup>a</sup> | 1.00 | 1.00 | 0.39 | 0.33 | 0.61 | 0.33 | 1.00 | 1.00 | 0.69 | 0.50 | 0.49 | 0.33 |
| Meta-SNP<sup>a</sup> | 1.00 | 1.00 | 0.68 | 0.35 | 0.88 | 0.38 | 1.00 | 1.00 | 0.91 | 0.54 | 0.78 | 0.37 |
| S3Ds&GO<sup>a</sup> | 1.00 | 1.00 | 0.53 | 0.24 | 0.86 | 0.36 | 1.00 | 1.00 | 0.88 | 0.47 | 0.68 | 0.29 |
| CADD | 0.96 | 0.95 | 1.00 | 0.73 | 1.00 | 0.90 | 0.85 | 0.85 | 0.97 | 0.88 | 0.90 | 0.71 |
| ConSurf | 0.98 | 0.97 | 0.82 | 0.50 | 0.96 | 0.77 | 0.90 | 0.90 | 0.95 | 0.80 | 0.83 | 0.56 |
| DANN | 0.93 | 0.89 | 0.90 | 0.60 | 0.98 | 0.85 | 0.69 | 0.69 | 0.92 | 0.81 | 0.75 | 0.51 |
| FATHMM | 1.00 | 1.00 | 0.52 | 0.45 | 0.76 | 0.59 | 1.00 | 1.00 | 0.81 | 0.69 | 0.63 | 0.51 |
| MAPP | 0.95 | 0.88 | 0.48 | 0.29 | 0.78 | 0.36 | 0.83 | 0.83 | 0.79 | 0.47 | 0.51 | 0.18 |
| MutPred2 | 1.00 | 1.00 | 0.72 | 0.36 | 0.90 | 0.41 | 1.00 | 1.00 | 0.92 | 0.56 | 0.81 | 0.38 |
| PANTHER-PSEP | 0.85 | 0.81 | nr | nr | 1.00 | 1.00 | 0.00 | 0.00 | 0.85 | 0.81 | nr | nr |
| PhD-SNPg | 0.98 | 0.95 | 0.57 | 0.39 | 0.82 | 0.51 | 0.92 | 0.92 | 0.84 | 0.62 | 0.64 | 0.38 |
| PolyPhen2-HumVar | 0.96 | 0.94 | 0.85 | 0.52 | 0.96 | 0.74 | 0.85 | 0.85 | 0.94 | 0.77 | 0.81 | 0.52 |
| PROVEAN | 1.00 | 1.00 | 0.76 | 0.39 | 0.92 | 0.49 | 1.00 | 1.00 | 0.94 | 0.62 | 0.84 | 0.44 |
| SIFT | 0.96 | 0.92 | 0.69 | 0.42 | 0.90 | 0.62 | 0.85 | 0.85 | 0.89 | 0.67 | 0.70 | 0.40 |
| SNP2 | 1.00 | 1.00 | 0.59 | 0.37 | 0.82 | 0.44 | 1.00 | 1.00 | 0.86 | 0.58 | 0.70 | 0.40 |
| SNPs&GO | 1.00 | 1.00 | 0.59 | 0.33 | 0.82 | 0.31 | 1.00 | 1.00 | 0.86 | 0.48 | 0.70 | 0.32 |

<sup>a</sup> meta-predictor. nr, no result; PPV, positive predictive value; NPV, negative predictive value; Se, sensibility; Sp, specificity; Ac, accuracy; MCC, Matthews' correlation coefficient test.

We obtained both sensibility and specificity higher than 0.8 for three predictors, CADD (se=0.96 and sp=0.85), ConSurf (se=0.88 and sp=0.90), and PolyPhen-2 (se=0.87 and sp=0.85). Moreover, five predictors obtained the accuracy between excellent and good, CADD (0.94), PANTHER-PSEP (0.91), DANN (0.89), ConSurf (0.88), and PolyPhen-2 (0.86) (Table).

The Matthews’ correlation coefficient (MCC) test showed positive values for that of almost all predictors (except by PANTHER-PSEP), being five of them with the MCC > 0.5 (Table). The best performance, value closer to +1, was obtained by CADD (0.75), followed by ConSurf (0.58), PolyPhen-2 (0.57), DANN (0.56), and PROVEN (0.51). Figure 3 shows a Venn diagram of the four predictors with better accuracy and MCC values.

**Figure 3.**
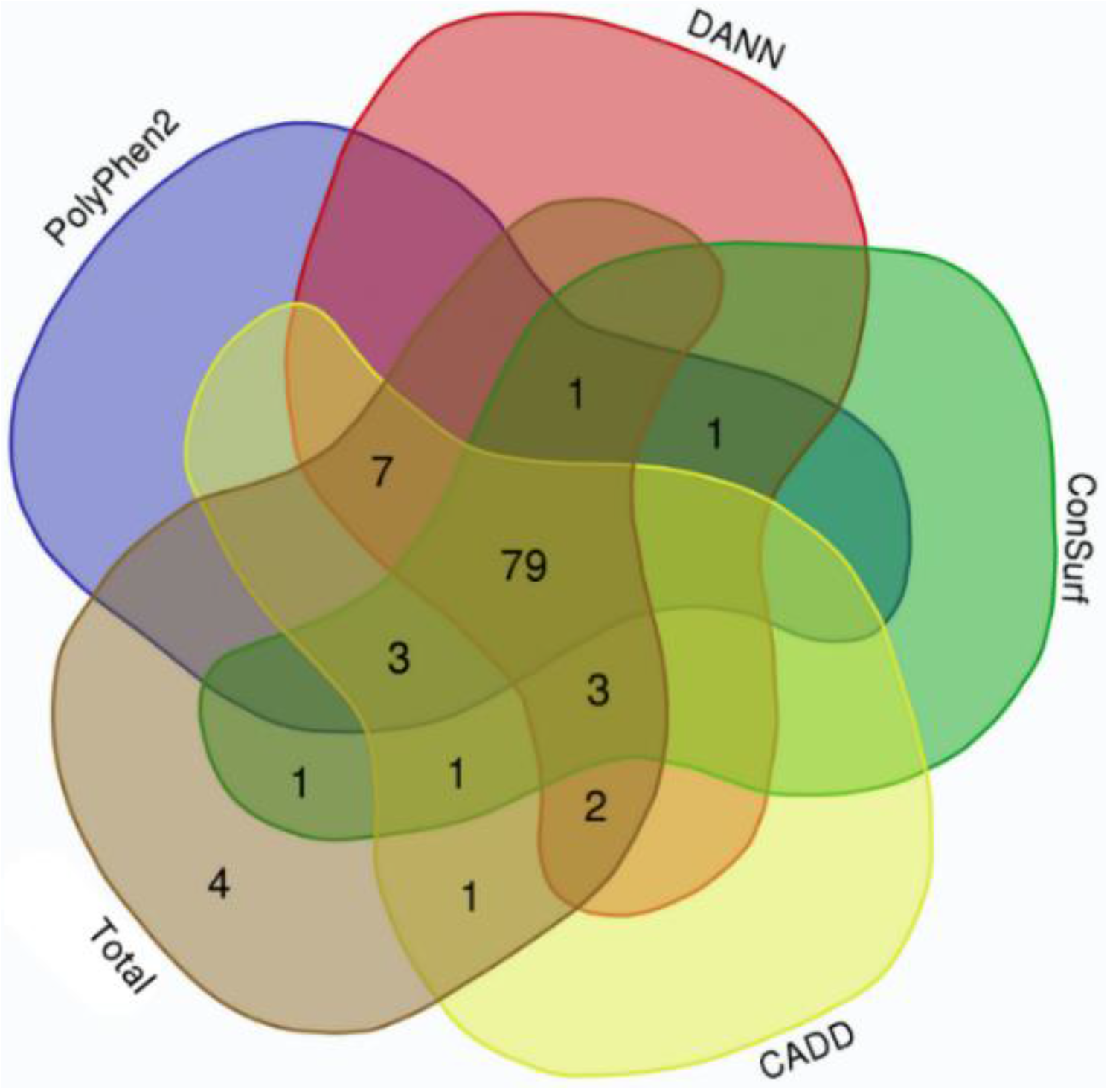
Venn diagram indicating overlaps of the hit for four predictors with good-excellent performance for SNVs on CYP21A2. Total, indicate all 103 SNVs tested. (Image generated with https://bioinformatics.psb.ugent.be/webtools/Venn/).

### Performance of predictors to identify SNVs affecting the specific CAH groups

We analyzed the performance of the selected predictors to identify 51 SNVs of the CL group and 39 SNVs of the NC group against 13 SNVs with a neutral effect (Table). Seventeen predictors obtained an excellent PPV rate (> 0.90) for the SNV CL group, 15 for the SNVs NC group. Four tools obtained excellent-good (> 0.8) NPV values for the CL group: CADD (1.0), DANN (0.9), PolyPhen-2 (0.85), and ConSurf (0.82). However, for the NC group, only CADD (0.73), DANN (0.6), and PolyPhen-2 (0.52) showed NPV > 0.5.

Taking the sensibility and specificity balance, we obtained 12 tools with excellent-good values (>0.8) for the CL group (Table). However, for the NC group, only CADD (se=0.9 and sp=0.85) obtained both sensibility and specificity with excellent-good values (Table). The accuracy was excellent for 7 predictors on CL group, CADD (0.97), ConSurf (0.95), PolyPhen-2 and PROVEN (0.94), DANN and MutPred2 (0.92), and Meta-SNP (0.91); and good for four tools on NC group, CADD (0.88), DANN and PANTHER-PSEP (0.81), and ConSurf (0.8).

Finally, the MCC test with positive values was obtained by almost all predictors (except for PANTHER-PSEP). The MCC was higher than 0.5 for 16 predictors in the CL group and five in the NC group (Table). For the CL and NC group, the same predictor, CADD, got the MCC value closer to +1, being MCC=0.9 for the CL group and MCC=0.71 for the NC group.

## Discussion

Genetics analysis is an essential approach for elucidating complex CYP21A2 deficiency cases, mainly to confirm asymptomatic carriers and unfollow false-positive cases ^7,16^. Therefore, fast and accessible tools to infer variants’ pathogenicity are essential to quickly deduce the harm of unknown variants. *In silico* prediction is one of the most accessible tools to infer the pathogenicity of SNVs.

Here, for the first time, we analyzed the performance of *in silico* prediction tools to discriminate pathogenic and neutral variants of the CYP21A2. We focus on the performance of 13 single predictors and four meta predictors chosen accordingly with the popularity and performance of free access programs. All these programs were able to identify pathogenic variants. Nonetheless, only PANTHER-PSEP could not distinguish neutral variants, which is unacceptable for testing variants of the CYP21A2. Moreover, all tools showed better performance with variants of the CL group than the NC group, as expected, since the CL group gathers the most harmful variants.

Our databank for performance tests comprises all missense variants of the CYP21A2 that are functionally characterized. With this strategy, we could get a more realistic result on the prediction evaluation. However, the number of variants was imbalanced between the two categories, 90 pathogenic and 13 neutral. Therefore, the primary statistics data considered for the performance evaluation were the accuracy and MCC, which consider all values (TP, TN, FP, and FN) ^34,35^. As sensibility and specificity are calculated with half of the information, they cannot represent all the performance by themselves, so we considered the sensibility-specificity balance. Additionally, we also calculated PPV and NPV but, as both are more sensitive to data disbalancing, they were not considered for the program performance ^34^.

The main feature assessed by most single predictors tested is the evolutionary data since residue conservation over time can indicate critical residues for the protein function. Four of the tools tested use only this feature for the prediction calculation, FATHMM ^20^, PhD-SNPg ^24^, PROVEAN ^26^, and SIFT ^27^. A similar performance was obtained between these four tools, with a fair accuracy ranging from 0.73 (FATHMM) to 0.79 (SIFT), and MCC from 0.42 (PhD-SNPg) to 0.51 (PROVEAN). Additionally, SIFT had the most excellent sensibility-specificity balance between programs, similar to the performance shown previously ^27^. In another study ^14^, with *HSD17B3, NR5A1, AR*, and *LHCGR* genes, SIFT and PROVEAN also had the same performance, with an accuracy of 0.74-0.75, and MCC of 0.5. In yet another study ^12^, SIFT and PROVEN showed the best results between nine programs tested for *GJB2, GJB6*, and *GJB3* genes, with an accuracy of 0.89, while FATHMM produced a large number of erroneous predictions with an accuracy of 0.33. FATHMM also had poor performance in another ^14^ study, with an accuracy of 0.56 and MCC of 0.04.

Changes in the secondary and tertiary structure by missense mutations are likely to affect the protein activity ^18^. Therefore, it is no surprise that the second most evaluated feature is the structural information, being present in ConSurf ^18^, MutPred2 ^22^, PolyPhen2 ^10^, and SNAP2 ^28^. In addition, ConSurf includes phylogenetics relationships, MutPred2 functional proprieties, and SNAP2 uses a matrix of effect probabilities with a neural network method ^18,22,28^. While, PolyPhen2 has two trained datasets as options, HumVar and HumDir. The first trained dataset is suggested for diagnostics of Mendelian diseases, which requires variants with a drastic difference effect ^25^. Moreover, PolyPhen2 has a low dependency on the sequence alignment employed ^10^. PolyPhen2 showed good prediction performance (ac=0.88, MCC=0.64), with the same sensibility but better sensibility-specificity balance than reported in ^10^. ConSurf obtained similar performance with the setting used in or test (ac=0.88, MCC=0.58). Since the ConSurf setting is chosen for the alignment sequences, databank, and algorithms, the performance is not comparable with other studies, as it was reported by ConSurf’s developers ^18^. On the other hand, MutPred2 and SNAP2 both had sensibility-specificity imbalanced of 1.4 and 1.5-folds, respectively, and almost the same fair performance. However, these values were relatively better than reported for MutPred2 by ^22^ with ClinVar and UniProt database and SNAP2 by ^28^ with a databank with more than 9,500 variants from human genes.

For meta predictors, which work with many databases and combine outputs from other predictors to generate their own, we expected to obtain one of the best performances. However, counterintuitively, our study showed an intermediate performance compared with the single predictors tested, and the number of tools combined was not related to the prediction improvement. Meta-SNP and PredictSNP performance were better than PredictSNP2 and S3Ds&GO. Furthermore, comparing with the developer tests, we obtained for Meta-SNP ^32^ and PredictSNP ^30^ a similar accuracy, while the performance for PredictSNP2 ^31^ and S3Ds&GO ^33^ were lower. Meta-SNP and PredictSNP share three single predictors, PhD-SNP, SNAP, and SIFT.

Therefore, for the CYP21A2 variants tested, the best performance to categorize missense variants pathogenicity was CADD, with an overall accuracy of 0.94 (CL 0.97; NC 0.88) and MCC of 0.75 (CL 0.9, NC 0.71). The specificity (0.85) and sensibility (0.9) also got a good balance. Interestingly, the accuracy and specificity obtained for CYP21A2 were even higher than reported by the software developers ^17^ with the ClinVar database, which was 0.85 and 0.57, respectively. ConSurf, DANN, and PolyPhen-2 showed a similar performance, holding the second-best results accordingly with the accuracy (CAH group 0.86-0.89; CL 0.92-0.95; NC 0.77-0.81,) and MCC (CAH group 0.56-0.58; CL 0.75-0.83; NC 0.51-0.56) values. The sensibility and specificity for ConSurf and PolyPhen2 were well balanced, while DANN had 1.3-folds less specificity than sensibility. The original article of DANN ^19^ presents only the area under the curve (AUC) ROC, which was 0.95 using the ClinVar database for the performance test. PolyPhen2 showed better sensibility-specificity balance for the CYP21A2 variants than the values presented by ^10^, testing the tool with gene-specific mutations (*BRCA1, MSH2, MLH1*, and *TP52*). We obtained a sensibility similar to a previous analysis ^10^, but the specificity was lower, 0.85 and 0.60, respectively.

The individual error of the top four predictors was six on CADD, 12 on ConSurf, 11 on DANN, and 14 on PolyPhen2. However, computing their prediction together, we would have four false results from 103 missense SNV in the CPY21A2, one neutral (p.L13M), and three pathogenic from the NC phenotype group (p.P106L, p.R225W, and p.M474I). The variant p.P106L was correctly categorized by SNAP2 and PANTHER-PSEP. In turn, PROVEAN^26^ could type the other three variants correctly, even with lower sensibility value than the other four tools, mainly for the pathogenic SNVs of the NC group (0.49). Nonetheless, we obtained an intermediated performance with PROVEAN (ac=0.77; MCC=0.51), which could be because it uses the neighborhood sequences as input, which can be a trick for enzymes since they have some residues with high conservation making critical connections between variable residues. Comparing with the developer’ test ^26^ with the UniProt database (se=0.78; sp=0.79), we had imbalanced sensibility-specificity imbalanced, with similar overall sensibility (0.73) and higher specificity (1.0).

In conclusion, we could identify CADD, ConSurf, DANN, and PolyPhen2 as good programs for missense variants prediction of the CYP21A2. Moreover, CADD had the best performance also for identifying mild mutations from the NC group, followed by ConSurf and DANN. These results may be applicable in future analysis of new or uncharacterized missense variants of the CYP21A2.

## Author Contributions

Conceptualization, M.J.P, R.L-B, A.Z., and A.V.P; methodology, M.J.P., A.V.P, and R.L-B; formal analysis, M.J.P; investigation, M.J.P.; resources, A.V.P. and A.Z.; writing—original draft preparation, M.J.P.; writing—review and editing, R.L-B., A.Z, L.M.R., and A.V.P.; visualization, M.J.P; supervision, A.V.P., A.Z., M.L.R.R., and R.L-B.; project administration, xx; funding acquisition, M.J.P., A.V.P., and A.Z. All authors have read and agreed to the published version of the manuscript.

## Supplementary Materials

Supplementary materials can be accessed at http://dx.doi.org/10.48350/162936 and https://boris.unibe.ch/id/eprint/162936

