## Supplementary materials for "Meta analysis of variant predictions in congenital adrenal hyperplasia caused by mutations in CYP21A2"

[1]Graduate Program in Cell and Molecular Biology, Universidade Federal do Rio Grande do Sul (UFRGS), Porto Alegre CEP 91501-970, Brazil. [2] Center for Biotechnology, Universidade Federal do Rio Grande do Sul (UFRGS), Porto Alegre CEP 91501-970, RS, Brazil. [3] Department of Biomedical Research, University of Bern, Bern 3010, Switzerland. [4] Pediatric Endocrinology Unit, Department of Pediatrics, University Children's Hospital Bern, Bern 3010, Switzerland. [5] Department of Pharmacosciences, Universidade Federal de Ciências da Saúde de Porto Alegre (UFCSPA), Porto Alegre CEP 90050-170, Brazil. [6] Graduate Program in Molecular Biology Applied to Health, Universidade Luterana do Brasil (ULBRA), Canoas CEP 92425-020, Brazil.

**ORCiD numbers:** M.J.P (0000-0003-0647-4429), R.L-B. (0000-0002-2555-9754), A.Z. (0000-0001-6336-474X), M.L.R.R (0000-0002-9672-9394), and A.V.P (0000-0001-8331-5902).

https://boris.unibe.ch/id/eprint/162936

DOI: http://dx.doi.org/10.48350/162936

Table S1. Single predictors selected for performance analysis with *CYP21A2* variants.

| Single Predictors | Description | Website | Ref. |
|---|---|---|---|
| CADD | Integrative annotation built based on diverse genomic feature derived from surrounding sequence context, gene model **annotation**, **evolutionary** constraint, **epigenetic** measurements, and **functional** predictions. | https://cadd.gs.washington.edu/ | [1] |
| ConSurf | Algorithm uses phylogenetic relationships among homologous sequences and the specific dynamics of the analyzed sequence with **evolutionary** models to estimate the evolutionary rates of the amino acid of the macromolecules and to map them onto the structure and/or sequence. | https://consurf.tau.ac.il/ | [2] |
| DANN | Deep neural network which takes non-linear relationships among features based on diverse genomic derived from surrounding sequence context, gene model **annotation**, **evolutionary** constraint, **epigenetic** measurements, and **functional** predictions. | https://cbcl.ics.uci.edu/public_data/DANN/ | [3] |
| FATHMM | **Evolutionary** conservation algorithm which uses homologous sequences with species-specific weighting to predict the protein's tolerance to missense variants. | http://fathmm.biocompute.org.uk/ | [4] |
| MAPP | A statistical framework predictor which uses protein **physicochemical** characteristics of each amino acid position on the **evolutionary** variation. | http://mendel.stanford.edu/sidowlab/downloads/MAPP/index.html | [5] |
| MutPred2 | Machine learning-based to predict amino acid substitution through **evolutionary**, **structural,** and **functional** proprieties. | http://mutpred.mutdb.org/ | [6] |
| PANTHER-PSEP | Predict using **evolutionary** preservation data, measuring though the length of time estimation that a site has been preserved. | http://www.pantherdb.org/tools/csnpScoreForm.jsp | [7] |
| PhD-SNP[g] | Machine learning algorithm for predicting SNVs in both non-coding and coding regions through **evolutionary data**. | https://snps.biofold.org/phd-snpg/ | [8] |
| PolyPhen-2 | It uses human protein **evolutionary** and **structural** data to predict amino acid substitution effect on the protein stability and functionality. | http://genetics.bwh.harvard.edu/pph2/ | [9] |
| PROVEN | Predict the functional effect through amino acid exchange **evolutionary** data and quality of the neighborhood sequence alignment rather than the target position. | http://provean.jcvi.org/genome_submit_2.php?species=human | [10] |
| SIFT | Predicts through sequence homology algorithm assuming **evolutionary** conserved regions tend to be less tolerant. | https://sift.bii.a-star.edu.sg/www/SIFT4G_vcf_submit.html | [11] |
| SNAP2 | A neural network method based on machine learning to predict the variant effect in the molecular function through **evolutionary** and **structural** protein data with an amino acid substitution matrix of effect probabilities. | https://rostlab.org/services/snap2web/ | [12] |
| SNPs&GO | Predict using **evolutionary** data, profile and gene ontology (biological process, cellular component and molecular function). When protein function is not available, it run PANTHER and PhD-SNP. | https://snps.biofold.org/snps-and-go/snps-and-go.html | [13] |

19

Table S2. Performance of predictor tools.

| Predictor | PPV | NPV | Se | Sp | Ac | MCC | AUC-ROC | Dataset | Ref. |
|---|---|---|---|---|---|---|---|---|---|
| CADD | | | 93.6 | 57.1 | 0.85 | | | ClinVar (2015) | [1] |
| DANN | | | | | | | 0.95 | ClinVar (2014) | [3] |
| FATHMM (weighted) | 0.85 | 0.8 | 0.78 | 0.87 | 0.82 | 0.65 | | SwissVar (2012) | [4] |
| MAPP | | | | | 0.626-0.767 | | | Experimental studies | [5] |
| Meta-SNP | 0.79 | 0.8 | 0.8 | 0.79 | 0.79 | 0.59 | 0.86 | SwissVar (2009-2012) | [14] |
| MutPred2 | 96 | | 42.3 | 95.6 | | | 84.9 | ClinVar32 (2015) and UniProt80 (2015) | [6] |
| PANTHER-PSEP | | | | | | | 0.721 | Derived from SwissVar | [7] |
| PhD-SNPg | 0.85 | 0.85 | 0.94 | 0.67 | 0.85 | 0.65 | 0.91 | NewClinvar (2016) | [8] |
| PolyPhen-2* | | | 0.85 | 0.6015 | | | 0.79 | Mutations on the genes BRCA1, MSH2, MLH1 and TP53 | [15] |
| PredictSNP | | | | | 0.642 | 0.281 | 0.7 | Protein Mutant Database (07Mar26) | [16] |
| PredictSNP2 | | | | | 0.773 | 0.55 | 0.804 | Mendelian diseases (multiple databases) | [17] |
| PROVEAN | | | 0.78 | 0.79 | | | | UniProt human protein | [10] |
| SNPs&GO | 0.83 | 0.8 | 0.78 | 0.85 | 0.82 | 0.63 | | Derived from Swiss-Prot (2008) | [13] |
| SIFT 4G | | | 0.8 | 0.735 | 0.7732 | 0.53 | | UniRef90 (2011) | [11] |
| SNAP2 | | | | | 0.688 | 0.24 | | Data set consisting of 9,657 variants from 678 human proteins | [12] |
| SNP&GO3d | 0.84 | 0.86 | 0.87 | 0.83 | 0.85 | 0.7 | 0.92 | Derived from Swiss-Prot (2009) | [18] |

*Data from an article recommended on the original developer article. PPV, positive predictive value; NPV, negative predictive value; Se, sensibility; Sp, specificity; Ac, accuracy; MCC, Matthews' correlation coefficient test.

Table S3..List of the 103 single nucleotide variants (SNVs) on *CYP21A2* gene selected to test the performance of predictor tools. SNVs are grouped into classical (enzyme activity < 10%), non-classical (between 10 and 78 %) and neutral (> 78 %) groups. The enzyme activity levels of both 21-hydroxylase substrates - 17-hydroxyprogesterone and progesterone - were obtained from the original paper of the functional characterization. The phenotype was obtained from either the same paper or the original description of the new SNV. [a] Shows the percentage of enzyme activity measured for the conversion of both 21-hydroxylase substrates, considering as 100 % the 21-hydroxylase wild type activity. 17OHP: 17-hydroxyprogesterone. SW: salt wasting. SV: simple-virilizing. NC: non-classical. ND: non-determinate.

| | | **CYP21A2 Activity *in vitro*** | | | | | |
|---|---|---|---|---|---|---|---|
| **Group** | **NP_000491.4** | **17OHP[a]** | **SD (±)** | **Progesterone** | **SD (±)** | **Phenotype** | **Publication** |
| CL | p.P31Q | 0.2 | 0.2 | 0 | 0 | SW | [19] |
| | p.G57R | 0.7 | ND | 1.4 | ND | SV | [20] |
| | p.G65E | 0 | ND | 0 | ND | SW | [21] |
| | p.I78T | 3 | 2 | 5 | 3 | SV | [22] |
| | p.G91V | 0 | ND | 0 | ND | SW | [23] |
| | p.L108R | 0.4 | ND | 0.3 | ND | SW | [20] |
| | p.S114F | 4 | 1 | 4 | 2 | SV | [24] |
| | p.L123P | 1.42 | 2.13 | -1.86 | 5.19 | SW | [25] |
| | p.V140E | 0.7 | 1.3 | 0.5 | 0.6 | SW | [26] |
| | p.L143P | 0.4 | | 0.4 | | SW | [20] |
| | p.C148R | 4.3 | 0.9 | 3.6ny | 1.8 | SV-NC | [26] |
| | p.L167P | 0.3 | 0.06 | 0.4 | 0.6 | SW | [27] |
| | p.L168P | 0.7 | ND | 0.4 | ND | SW | [28] |
| | p.C170R | 0.1 | 0.02 | 0 | 2 | SW | [29] |
| | p.I172N | 0.7 | 0.3 | 0.6 | 0.03 | SV | [30] |
| | p.I173N | 4.3 | 1.7 | 4.4 | 1.8 | SV | [28] |
| | p.G179R | 0.4 | 0.5 | 0 | 0.6 | SW | [29] |
| | p.R234G | 8 | 2 | 2 | 1 | SV-NC | [31] |
| | p.I237N | 1 | 0.7 | 2.4 | 1.4 | SV | [32] |
| | p.V238E | 0 | 0 | 0.1 | 0.3 | SW | [32] |
| | p.V282G | 3.9 | 1.7 | 3.9 | 2 | SV | [33] |
| | p.H283N | 1.6 | 6 | 2.7 | 5 | SW | [34] |
| | p.G292C | 0 | ND | 0 | ND | SW | [23] |
| | p.G292R | 0.5 | 0.7 | 0.7 | 0.2 | SW | [26] |
| | p.G292S | 0.8 | 0.4 | 0.8 | 0.4 | SW | [35] |
| | p.G293D | 0.5 | 0.2 | 0.7 | 0.4 | SW | [28] |
| | p.L301F | 9.5 | 6.4 | 4.4 | 2.5 | SV | [33] |
| | p.W303S | 3 | 0.3 | 3 | 0.5 | SV-NC | [36] |
| | p.W303R | 0.1 | 0.2 | 0 | 0.5 | SW | [29] |
| | p.L309F | 0.2 | 0.3 | 0.1 | 0.3 | SW | [26] |
| | p.E321K | 4.6 | 1.8 | 4.5 | 2.6 | SV | [28] |
| | p.R342P | 0.7 | 0.3 | 0.7 | 0.2 | SV | [30] |
| | p.R342W | 5 | 0.4 | 4 | 3 | SV-NC | [31] |
| | p.E352K | 1.1 | 0.5 | 1.2 | 0.3 | SV | [37] |
| | p.R355H | 0 | ND | 0 | ND | SW | [23] |
| | p.R357P | 0.15 | 0.3 | 0.15 | 0.3 | SW | [38] |
| | p.R357Q | 0.65 | 0.44 | 1.1 | 0.94 | SV | [38] |
| | p.R357W | 0 | ND | 0 | ND | SW | [39] |
| | p.A363V | 0 | ND | 0 | ND | SW | [21] |
| | p.G376S | 1.6 | 0.8 | 0.7 | 0.7 | SW | [40] |
| | p.L389R | 1.1 | 0.6 | ND | ND | SW | [41] |

Continuation (Table S3)

| | | | | | | | |
|---|---|---|---|---|---|---|---|
| CL | p.H393Q | 2.5 | 0.6 | 2.2 | 0.6 | SW | [42] |
| | p.R409C | 1.3 | 0.5 | | | SW | [20] |
| | p.G425S | 1.6 | 0.4 | 2 | 0.6 | SV | [28] |
| | p.R427C | 0 | 0.5 | 0 | 0.6 | SW | [29] |
| | p.R427H | 0.5 | 0.6 | 0.4 | 0.2 | SW-SV | [30] |
| | p.L447P | 0.5 | 0.6 | 0 | 0.1 | SW-SV | [30] |
| | p.T451P | 0.9 | ND | 0.9 | ND | SW | [24] |
| | p.P464L | 2.6 | 0.8 | 3 | 0.5 | SV | [43] |
| | p.R484P | 1 | 0.07 | 2.2 | 0.9 | SV | [35] |
| | p.R484Q | 1.1 | 0.7 | 3.8 | 1.9 | SV | [27] |
| | **Mean** | **1.52** | | **1.32** | | | |
| | **SD** | **2.00** | | **1.61** | | | |
| NC | p.P31L | 13 | 0.2 | 2 | 0.6 | NC | [31] |
| | p.H63L | 44.5 | ND | 20.7 | ND | NC | [20] |
| | p.P106L | 62 | 9 | 64 | 12 | NC | [44] |
| | p.H120R | 31.6 | 8 | 32.5 | 7 | NC | [45] |
| | p.K122Q | 14 | 5 | 19.5 | 4 | NC | [46] |
| | p.R133C | 35.4 | 7.4 | 15.5 | 2.7 | NC | [47] |
| | p.E141K | 11.3 | 2.4 | ND | ND | SW | [41] |
| | p.R150C | 35.8 | 14.6 | 47.3 | 12.9 | NC | [47] |
| | p.R150P | 23.4 | 1.7 | 16.9 | 2 | NC | [48] |
| | p.M151R | 17.66 | 1.87 | 4.57 | 1.96 | NC | [25] |
| | p.G179A | 19 | ND | ND | ND | NC | [23] |
| | p.Y192H | 37.1 | 7 | 25.8 | 9 | NC | [34] |
| | p.I195N | 33.2 | 9 | 46.7 | 10 | NC | [45] |
| | p.R225W | 51.9 | 9 | 45.6 | 8 | NC | [49] |
| | p.I231T | 63.1 | 22.3 | 70.6 | 17 | NC | [28] |
| | p.R234K | 15 | ND | 8.1 | ND | SV-NC | [28] |
| | p.V282L | 18 | 3 | 18 | 5 | NC | [31] |
| | p.M284V | 16.2 | 9.3 | 19 | 6.8 | NC | [47] |
| | p.V305M | 46 | 18 | 26 | 10 | NC | [50] |
| | p.F307V | 63.23 | 5.5 | 64.17 | 7.98 | SV-NC | [51] |
| | p.D323G | 18 | 1.2 | 27 | 4.7 | NC | [36] |
| | p.R340H | 67.1 | 2.4 | 45.8 | 3.7 | NC | [52] |
| | p.V359I | 72 | 7 | 34 | 3 | NC | [53] |
| | p.H366N | 46.13 | 4.8 | 57.77 | 3.69 | NC | [51] |
| | p.R367C | 37 | 7 | 28 | 4 | NC | [31] |
| | p.R370Q | 82 | 6 | 63 | 4 | NC | [53] |
| | p.R370W | 45.8 | 1.8 | 48.5 | 17.1 | NC | [28] |
| | p.D378Y | 81 | 6 | 58 | 4 | NC | [53] |
| | p.E381D | 30 | ND | ND | ND | SW | [54] |
| | p.A392T | 38.7 | 9.5 | 22.9 | 4.7 | NC | [55] |
| | p.D408N | 72.7 | 7 | 73.6 | 10 | NC | [49] |
| | p.E432K | 26.2 | 3.8 | 24.2 | 7.4 | NC | [47] |
| | p.A435V | 14 | 2 | 12 | 6 | SV | [22] |
| | p.T451M | 78 | 6 | 43 | 5 | NC | [24] |
| | p.P454S | 38 | ND | 22.4 | 3 | NC | [31] |
| | p.L462P | 55 | 8 | 40 | 2 | NC | [53] |
| | p.M474I | 85 | 7 | 66 | 12 | NC | [31] |
| | p.R480L | 75.5 | 15.7 | 79.6 | 12 | NC-Normal | [55] |
| | p.P483S | 61 | 6 | 54 | 2 | NC | [31] |
| | **Mean** | **42.94** | | **37.41** | | | |
| | **SD** | **22.59** | | **20.98** | | | |

Continuation (Table S3)

| | | | | | | | |
|---|---|---|---|---|---|---|---|
| Neutral | p.L13M | 99 | 1 | 100 | 1 | Normal | [24] |
| | p.A16T | 100 | 0 | 96 | 6 | Normal- very mildNC | [24] |
| | p.R17C | 95 | 3 | 81 | 3 | Normal- very mildNC | [24] |
| | p.R103K | 119.7 | 22.5 | ND | ND | Normal | [41] |
| | p.A160T | 126.6 | 29.9 | ND | ND | Normal | [41] |
| | p.D184E | 100 | ND | 100 | ND | Normal | [56] |
| | p.S203G | 85 | 2 | 81 | 3 | Very mild NC | [24] |
| | p.V212M | 99.5 | 32.4 | ND | ND | Normal | [41] |
| | p.M240K | 95.4 | 24.7 | 97.7 | 7.7 | Normal | [32] |
| | p.A266S | 90 | 9 | 104 | 15 | Normal | [31] |
| | p.A266V | 92 | 1.4 | 100 | 4.3 | Normal | [36] |
| | p.P268L | 97 | 1 | 87 | 7 | Normal | [24] |
| | p.S269T | 103 | 15 | ND | ND | Normal | [57] |
| | **Average** | **100.17** | | **94.08** | | | |
| | **SD** | **10.92** | | **8.25** | | | |

Table S4. Result of 17 predictors for 51 classical single nucleotide variants (SNVs) on the CYP21A2 gene. The classical group has an enzyme activity of < 10% of the wild-type activity. The genomic SNV nomenclature is based on the human chromatin remodeling 38 (Chr38). Del: deleterious; N: Neutral; Pby: Probably; Psb: Possible; B: Benign; Dse: Disease; Efc: Effect; P-Del: Proxy-deleterious; P-N: Proxy-neutral; Dmg: Damaging; T: Tolerated; Ptg: Pathogenic; Csv: Conserved; V: Variable; NR: no result.

| Chr38 | SNP | Meta-SNP | PredictSNP | PredictSNP2 | S3Ds&GO | CADD | ConSurf | DANN | FATHMM | MAPP | MutPred2 | PANTHER | PhD-SNPg | PolyPhen2 | PROVEAN | SIFT | SNAP2 | SNPs&GO |
|---|---|---|---|---|---|---|---|---|---|---|---|---|---|---|---|---|---|---|
| g.32038514C>A | p.P31Q | Dse | Del | N | Dse | P-Del | Csv | Del | Dmg | Del | Del | Pby | Ptg | Pby | Del | Del | Efc | N; |
| g.32038591G>A | p.G57R | Dse | Del | Del | Dse | P-Del | Csv | Del | Dmg | Del | Del | Pby | Ptg | Pby | Del | Del | Efc | Dse |
| g.32038616G>A | p.G65E | Dse | Del | Del | Dse | P-Del | Csv | Del | Dmg | Del | Del | Pby | Ptg | Pby | Del | Del | Efc | Dse |
| g.32038752T>C | p.I78T | N | Del | Del | Dse | P-Del | Csv | Del | Dmg | Del | Del | Pby | Ptg | Pby | N | N | N | N |
| g.32038791G>T | p.G91V | Dse | Del | Del | Dse | P-Del | Csv | Del | Dmg | Del | Del | Pby | Ptg | Pby | Del | Del | Efc | Dse |
| g.32039124T>G | p.L108R | Dse | Del | N | Dse | P-Del | Csv | Del | Dmg | Del | Del | Pby | Ptg | Pby | Del | Del | Efc | Dse |
| g.32039142C>T | p.S114F | Dse | Del | Del | Dse | P-Del | Csv | Del | Dmg | Del | Del | Pby | B | Pby | Del | Del | Efc | Dse |
| g.32039169T>C | p.L123P | Dse | Del | N | Dse | P-Del | Csv | Del | T | Del | Del | Pby | Ptg | Pby | Del | Del | Efc | Dse |
| g.32039220T>A | p.V140E | Dse | Del | N | Dse | P-Del | Csv | Del | T | Del | Del | Pby | Ptg | Pby | Del | Del | Efc | Dse |
| g.32039229T>C | p.L143P | Dse | Del | N | Dse | P-Del | Csv | Del | T | Del | Del | Pby | Ptg | Pby | N | N | N | Dse |
| g.32039243T>C | p.C148R | N | N | N | N | P-Del | Csv | N | Dmg | Del | Del | Pby | Ptg | Psb | Del | N | N | N |
| g.32039408T>C | p.L167P | Dse | Del | N | Dse | P-Del | V | Del | T | Del | Del | Pby | Ptg | Pby | Del | Del | Efc | Dse |
| g.32039411T>C | p.L168P | Dse | Del | Del | Dse | P-Del | Csv | Del | Dmg | Del | Del | Pby | B | Pby | N | N | N | Dse |
| g.32039416T>C | p.C170R | Dse | Del | N | Dse | P-Del | Csv | Del | Dmg | Del | Del | Pby | B | Pby | Del | Del | Efc | Dse |
| g.32039423T>A | p.I172N | Dse | Del | N | Dse | P-Del | Csv | Del | Dmg | Del | Del | Pby | Ptg | Pby | Del | Del | Efc | Dse |
| g.32039426T>A | p.I173N | Dse | Del | Del | Dse | P-Del | Csv | Del | Dmg | Del | Del | Pby | Ptg | Pby | Del | Del | Efc | Dse |

Continuation (Table S4)

| Chr38 | SNP | Meta-SNP | PredictSNP | PredictSNP2 | S3Ds&GO | CADD | ConSurf | DANN | FATHMM | MAPP | MutPred2 | PANTHER | PhD-SNPg | PolyPhen2 | PROVEAN | SIFT | SNAP2 | SNPs&GO |
|---|---|---|---|---|---|---|---|---|---|---|---|---|---|---|---|---|---|---|
| g.32039443G>A | p.G179R | Dse | Del | Del | Dse | P-Del | Csv | Del | Dmg | Del | Del | Pby | Ptg | Pby | Del | Del | Efc | Dse |
| g.32039797A>G | p.R234G | Dse | N | N | Dse | P-Del | Csv | Del | T | N | Del | Pby | B | Pby | Del | Del | Efc | Dse |
| g.32039807T>A | p.I237N | Dse | Del | N | N | P-Del | V | Del | T | Del | Del | Pby | Ptg | Psb | Del | Del | Efc | Dse |
| g.32039810T>A | p.V238E | Dse | Del | N | Dse | P-Del | Csv | Del | T | Del | Del | Pby | Ptg | Psb | Del | Del | Efc | Dse |
| g.32040111T>G | p.V282G | Dse | Del | Del | Dse | P-Del | Csv | Del | Dmg | Del | Del | Pby | Ptg | Pby | Del | Del | Efc | Dse |
| g.32040113C>A | p.H283N | Dse | N | N | Dse | P-Del | Csv | Del | Dmg | N | N | Pby | Ptg | Pby | Del | Del | N | Dse |
| g.32040140G>A | p.G292S | Dse | Del | Del | Dse | P-Del | Csv | Del | Dmg | Del | Del | Pby | Ptg | Pby | Del | Del | Efc | Dse |
| g.32040140G>C | p.G292R | Dse | Del | Del | Dse | P-Del | Csv | Del | Dmg | Del | Del | Pby | Ptg | Pby | Del | Del | Efc | Dse |
| g.32040140G>T | p.G292C | Dse | Del | Del | Dse | P-Del | Csv | Del | Dmg | Del | Del | Pby | Ptg | Pby | Del | Del | Efc | Dse |
| g.32040144G>A | p.G293D | Dse | Del | Del | Dse | P-Del | Csv | Del | Dmg | Del | Del | Pby | Ptg | Pby | Del | Del | Efc | Dse |
| g.32040167C>T | p.L301F | Dse | Del | Del | Dse | P-Del | Csv | Del | Dmg | N | Del | Pby | B | Pby | Del | Del | Efc | Dse |
| g.32040173T>C | p.W303R | Dse | Del | Del | Dse | P-Del | Csv | Del | Dmg | Del | Del | Pby | Ptg | Pby | Del | Del | Efc | Dse |
| g.32040174G>C | p.W303S | Dse | Del | Del | Dse | P-Del | Csv | Del | Dmg | Del | Del | Pby | Ptg | Pby | Del | Del | Efc | Dse |
| g.32040191C>T | p.L309F | N | N | Del | N | P-Del | Csv | Del | Dmg | N | N | Pby | B | Pby | Del | Del | N | N |
| g.32040427G>A | p.E321K | Dse | Del | Del | Dse | P-Del | Csv | Del | Dmg | Del | Del | Pby | Ptg | Pby | Del | Del | Efc | Dse |
| g.32040490C>T | p.R342W | Dse | Del | N | Dse | P-Del | Csv | Del | Dmg | N | Del | Pby | Ptg | Pby | Del | Del | Efc | Dse |
| g.32040491G>C | p.R342P | Dse | Del | N | Dse | P-Del | Csv | Del | T | Del | Del | Pby | B | Pby | Del | Del | Efc | Dse |
| g.32040520G>A | p.E352K | Dse | Del | Del | Dse | P-Del | Csv | Del | Dmg | Del | Del | Pby | Ptg | Pby | Del | Del | Efc | Dse |
| g.32040530G>A | p.R355H | Dse | Del | Del | Dse | P-Del | Csv | Del | Dmg | Del | Del | Pby | Ptg | Psb | Del | Del | Efc | Dse |

Continuation (Table S4)

| Chr38 | SNP | Meta-SNP | PredictSNP | PredictSNP2 | S3Ds&GO | CADD | ConSurf | DANN | FATHMM | MAPP | MutPred2 | PANTHER | PhD-SNPg | PolyPhen2 | PROVEAN | SIFT | SNAP2 | SNPs&GO |
|---|---|---|---|---|---|---|---|---|---|---|---|---|---|---|---|---|---|---|
| g.32040535C>T | p.R357W | Dse | Del | N | Dse | P-Del | Csv | Del | T | N | Del | Pby | Ptg | Pby | Del | Del | Efc | Dse |
| g.32040536G>A | p.R357Q | Dse | Del | Del | Dse | P-Del | Csv | Del | Dmg | N | N | Pby | Ptg | Pby | Del | Del | Efc | Dse |
| g.32040536G>C | p.R357P | Dse | Del | Del | Dse | P-Del | Csv | Del | Dmg | Del | Del | Pby | Ptg | Pby | Del | Del | Efc | Dse |
| g.32040554C>T | p.A363V | N | N | N | Dse | P-Del | Csv | Del | T | N | N | Pby | Ptg | Pby | N | N | N | N |
| g.32040675G>A | p.G376S | Dse | Del | Del | Dse | P-Del | Csv | Del | Dmg | N | Del | Pby | Ptg | Pby | Del | Del | Efc | Dse |
| g.32040715T>G | p.L389R | Dse | Del | Del | Dse | P-Del | Csv | Del | Dmg | Del | Del | Pby | Ptg | Pby | Del | Del | Efc | Dse |
| g.32040728C>G | p.H393Q | N | N | N | Dse | P-Del | Csv | Del | T | N | Del | Pby | B | B | Del | Del | Efc | N |
| g.32040871C>T | p.R409C | Dse | Del | Del | Dse | P-Del | Csv | Del | Dmg | Del | Del | Pby | Ptg | Psb | Del | Del | Efc | Dse |
| g.32040919G>A | p.G425S | Dse | Del | Del | Dse | P-Del | Csv | Del | Dmg | Del | Del | Pby | Ptg | Pby | Del | Del | Efc | Dse |
| g.32040925C>T | p.R427C | Dse | Del | Del | Dse | P-Del | Csv | Del | Dmg | Del | Del | Pby | Ptg | Pby | Del | Del | Efc | Dse |
| g.32040926G>A | p.R427H | Dse | Del | Del | Dse | P-Del | Csv | Del | Dmg | Del | Del | Pby | Ptg | Pby | Del | Del | Efc | Dse |
| g.32040986T>C | p.L447P | Dse | Del | Del | Dse | P-Del | Csv | Del | Dmg | Del | Del | Pby | B | Pby | Del | Del | Efc | Dse |
| g.32040997A>C | p.T451P | Dse | Del | N | N | P-Del | Csv | Del | T | Del | Del | Pby | Ptg | Psb | Del | Del | N | N |
| g.32041037C>T | p.P464L | N | N | Del | N | P-Del | Csv | Del | Dmg | Del | N | Pby | Ptg | Pby | Del | Del | N | N |
| g.32041097G>A | p.R484Q | Dse | Del | Del | N | P-Del | Csv | Del | Dmg | N | Del | Pby | Ptg | Pby | Del | Del | Efc | N |
| g.32041097G>C | p.R484P | Dse | Del | N | N | P-Del | Csv | Del | Dmg | Del | Del | Pby | Ptg | Pby | Del | Del | Efc | Dse |

Table S5. Result of 17 predictors for 39 non-classical single nucleotide variants (SNVs) on the CYP21A2. The non-classical group has an enzyme activity between >10% and < 78% of the wild-type activity. The genomic SNV nomenclature is based on the human chromatin remodeling 38 (Chr38). Del: deleterious; N: Neutral; Pby: Probably; Psb: Possible; B: Benign; Dse: Disease; Efc: Effect; P-Del: Proxy-deleterious; P-N: Proxy-neutral; Dmg: Damaging; T: Tolerated; Ptg: Pathogenic; Csv: Conserved; V: Variable; NR: no result.

| Chr38 | SNP | Meta-SNP | PredictSNP | PredictSNP2 | S3Ds&GO | CADD | ConSurf | DANN | FATHMM | MAPP | MutPred2 | PANTHER | PhD-SNPg | PolyPhen2 | PROVEAN | SIFT | SNP2 | SNPs&GO |
|---|---|---|---|---|---|---|---|---|---|---|---|---|---|---|---|---|---|---|
| g.32038514C>T | p.P31L | N | N | N | N | P-Del | Csv | N | Dmg | Del | Del | Pby | Ptg | B | N | N | N | N |
| g.32038610A>T | p.H63L | Dse | N | N | N | P-Del | V | N | T | Del | Del | Pby | B | B | N | N | Efc | N |
| g.32039118C>T | p.P106L | N | N | N | N | P-N | V | N | T | N | N | Pby | B | B | N | N | Efc | N |
| g.32039160A>G | p.H120R | N | N | N | N | P-Del | Csv | Del | Dmg | Del | N | Pby | Ptg | Pby | Del | Del | Efc | N |
| g.32039165A>C | p.K122Q | Dse | Del | N | N | P-Del | Csv | Del | Dmg | Del | N | Pby | Ptg | Pby | Del | Del | Efc | N |
| g.32039198C>T | p.R133C | Dse | Del | N | Dse | P-Del | V | Del | T | N | N | Pby | Ptg | Pby | Del | Del | Efc | Dse |
| g.32039222G>A | p.E141K | N | N | N | N | P-Del | V | Del | T | N | Del | Pby | Ptg | Psb | N | N | N | N |
| g.32039356C>T | p.R150C | N | N | Del | Dse | P-Del | Csv | Del | Dmg | N | N | Pby | B | Pby | N | N | N | N |
| g.32039357G>C | p.R150P | Dse | Del | N | Dse | P-Del | Csv | Del | Dmg | Del | Del | Pby | Ptg | Pby | N | N | N | Dse |
| g.32039360T>G | p.M151R | Dse | Del | N | Dse | P-Del | Csv | Del | Dmg | Del | Del | Pby | Ptg | Psb | Del | Del | Efc | Dse |
| g.32039444G>C | p.G179A | Dse | Del | Del | Dse | P-Del | Csv | Del | Dmg | Del | N | Pby | Ptg | Pby | Del | Del | Efc | Dse |
| g.32039570T>C | p.Y192H | N | N | N | N | P-N | Csv | N | T | N | N | Pby | B | B | N | N | N | N |
| g.32039580T>A | p.I195N | N | N | N | Dse | P-Del | Csv | Del | T | N | Del | Pby | Ptg | Pby | N | Del | Efc | Dse |
| g.32039770C>T | p.R225W | Dse | N | N | N | P-N | V | N | T | N | Del | Pby | B | B | Del | N | N | N |
| g.32039789T>C | p.I231T | N | N | N | N | P-Del | V | Del | T | N | N | Pby | B | B | N | Del | N | N |
| g.32039798G>A | p.R234K | N | N | N | Dse | P-Del | Csv | Del | Dmg | N | N | Pby | Ptg | Pby | N | Del | Efc | N |

Continuation (Table S5)

| Chr38 | SNP | Meta-SNP | PredictSNP | PredictSNP2 | S3Ds&GO | CADD | ConSurf | DANN | FATHMM | MAPP | MutPred2 | PANTHER | PhD-SNPg | PolyPhen2 | PROVEAN | SIFT | SNP2 | SNPs&GO |
|---|---|---|---|---|---|---|---|---|---|---|---|---|---|---|---|---|---|---|
| g.32040110G>T | p.V282L | N | N | N | N | P-Del | Csv | Del | Dmg | N | N | Pby | Ptg | Psb | N | N | N | N |
| g.32040116A>G | p.M284V | N | N | Del | Dse | P-Del | Csv | Del | Dmg | N | N | Pby | Ptg | Pby | Del | Del | Efc | Dse |
| g.32040179G>A | p.V305M | Dse | N | Del | N | P-Del | Csv | Del | Dmg | N | N | Pby | B | Pby | N | Del | N | N |
| g.32040185T>G | p.F307V | N | N | N | Dse | P-Del | Csv | Del | Dmg | N | Del | Pby | B | Pby | Del | Del | N | Dse |
| g.32040434A>G | p.D323G | Dse | Del | N | N | P-Del | V | Del | T | Del | Del | Pby | B | Pby | Del | Del | N | N |
| g.32040485G>A | p.R340H | Dse | Del | Del | N | P-Del | Csv | Del | Dmg | N | Del | Pby | B | Pby | Del | Del | Efc | Dse |
| g.32040541G>A | p.V359I | N | N | N | N | P-Del | Csv | Del | Dmg | N | N | Pby | B | Psb | N | Del | N | N |
| g.32040562C>A | p.H366N | Dse | Del | N | Dse | P-Del | Csv | Del | Dmg | Del | Del | Pby | Ptg | Pby | Del | Del | Efc | N |
| g.32040565C>T | p.R367C | N | N | N | N | P-Del | Csv | Del | T | N | Del | Pby | B | B | N | N | N | N |
| g.32040574C>T | p.R370W | Dse | Del | N | Dse | P-Del | Csv | Del | T | Del | N | Pby | B | Pby | Del | Del | Efc | Dse |
| g.32040575G>A | p.R370Q | N | N | N | N | P-Del | Csv | Del | T | N | N | Pby | B | Psb | N | N | N | N |
| g.32040681G>T | p.D378Y | N | N | N | N | P-Del | Csv | Del | T | N | Del | Pby | B | Psb | Del | N | N | N |
| g.32040692G>C | p.E381D | N | N | Del | N | P-Del | Csv | Del | Dmg | N | N | Pby | B | B | N | Del | Efc | N |
| g.32040723G>A | p.A392T | N | N | Del | Dse | P-Del | Csv | Del | Dmg | Del | N | Pby | B | Pby | N | N | N | Dse |
| g.32040771G>A | p.D408N | N | N | Del | N | P-Del | Csv | Del | Dmg | N | N | Pby | Ptg | Pby | N | Del | N | N |
| g.32040940G>A | p.E432K | Dse | Del | Del | Dse | P-Del | Csv | Del | Dmg | N | Del | Pby | Ptg | Pby | Del | Del | Efc | Dse |
| g.32040950C>T | p.A435V | Dse | Del | Del | Dse | P-Del | Csv | Del | Dmg | Del | Del | Pby | Ptg | Pby | Del | Del | Efc | Dse |
| g.32040998C>T | p.T451M | N | Del | N | N | P-Del | Csv | Del | T | Del | N | Pby | Ptg | Psb | Del | Del | N | N |
| g.32041006C>T | p.P454S | N | Del | Del | N | P-Del | Csv | Del | Dmg | N | N | Pby | Ptg | Pby | Del | Del | N | N |

Continuation (Table S5)

| **Chr38** | **SNP** | **Meta-SNP** | **PredictSNP** | **PredictSNP2** | **S3Ds&GO** | **CADD** | **ConSurf** | **DANN** | **FATHMM** | **MAPP** | **MutPred2** | **PANTHER** | **PhD-SNPg** | **PolyPhen2** | **PROVEAN** | **SIFT** | **SNP2** | **SNPs&GO** |
|---|---|---|---|---|---|---|---|---|---|---|---|---|---|---|---|---|---|---|
| g.32041031T>C | p.L462P | Dse | Del | Del | N | P-Del | Csv | Del | Dmg | Del | Del | Pby | Ptg | Pby | Del | Del | N | N |
| g.32041068G>T | p.M474I | N | N | N | N | P-N | V | N | T | N | N | Pby | B | B | Del | N | N | N |
| g.32041085G>T | p.R480L | N | N | N | N | P-Del | V | Del | T | N | N | Pby | B | B | N | N | N | N |
| g.32041093C>T | p.P483S | N | N | Del | N | P-Del | Csv | Del | Dmg | N | N | Pby | Ptg | Pby | N | Del | Efc | N |

Table S6. Result of 17 predictors for 13 neutral single nucleotide variants (SNVs) on the CYP21A2 gene. The neutral group has the enzyme activity known as > 78% of the wild-type activity. The genomic SNV nomenclature is based on the human chromatin remodeling 38 (Chr38). Del: deleterious; N: Neutral; Pby: Probably; Psb: Possible; Dse: Disease; Efc: Effect; P-Del: Proxy-deleterious; P-N: Proxy-neutral; Dmg: Damaging; T: Tolerated; Ptg: Pathogenic; B: Benign; Csv: Conserved; V: Variable; NR: no result.

| Chr38 | SNP | Meta-SNP | PredictSNP | PredictSNP2 | S3Ds&GO | CADD | ConSurf | DANN | FATHMM | MAPP | MutPred2 | PANTHER | PhD-SNPg | PolyPhen2 | PROVEAN | SIFT | SNP2 | SNPs&GO |
|---|---|---|---|---|---|---|---|---|---|---|---|---|---|---|---|---|---|---|
| g.32038459C>A | p.L13M | N | N | N | NR | P-Del | NR | Del | T | Del | N | NR | B | Pby | N | Del | N | N |
| g.32038468G>A | p.A16T | N | N | N | NR | P-N | NR | N | T | N | N | NR | B | B | N | N | N | N |
| g.32038471C>T | p.R17C | N | N | N | NR | P-N | NR | Del | T | N | N | NR | B | B | N | Del | N | N |
| g.32039109G>A | p.R103K | N | N | N | N | P-N | V | N | T | N | N | NR | B | B | N | N | N | N |
| g.32039386G>A | p.A160T | N | N | N | N | P-N | Csv | N | T | N | N | Pby | B | B | N | N | N | N |
| g.32039548C>G | p.D184E | N | N | N | N | P-N | V | N | T | N | N | Pby | B | B | N | N | N | N |
| g.32039603A>G | p.S203G | N | N | N | N | P-N | V | N | T | N | N | Pby | B | B | N | N | N | N |
| g.32039630G>A | p.V212M | N | N | N | N | P-N | V | Del | T | N | N | Pby | B | B | N | N | N | N |
| g.32039816T>A | p.M240K | N | N | N | N | P-Del | V | N | T | N | N | Pby | Ptg | B | N | N | N | N |
| g.32040062G>T | p.A266S | N | N | N | N | P-N | V | N | T | N | N | Pby | B | B | N | N | N | N |
| g.32040063C>T | p.A266V | N | N | N | N | P-N | V | Del | T | Del | N | Pby | B | B | N | N | N | N |
| g.32040069C>T | p.P268L | N | N | N | NR | P-N | V | N | T | N | N | Pby | B | B | N | N | N | N |
| g.32040072G>C | p.S269T | N | N | N | NR | P-N | V | N | T | NR | N | Pby | B | B | N | N | N | N |

35. Nikoshkov, A.; Lajic, S.; Vlamis-Gardikas, A.; Tranebjærg, L.; Holst, M.; Wedell, A.; Luthman, H. Naturally Occurring Mutants of Human Steroid 21-Hydroxylase (P450c21)

Pinpoint Residues Important for Enzyme Activity and Stability. *Journal of Biological Chemistry* **1998**, *273*, 6163–6165, doi:10.1074/jbc.273.11.6163.

46. Riepe, F.G.; Hiort, O.; Grötzinger, J.; Sippell, W.G.; Krone, N.; Holterhus, P.-M. Functional and Structural Consequences of a Novel Point Mutation in the *CYP21A2* Gene Causing Congenital Adrenal Hyperplasia: Potential Relevance of Helix C for P450

Oxidoreductase-21-Hydroxylase Interaction. *The Journal of Clinical Endocrinology & Metabolism* **2008**, *93*, 2891–2895, doi:10.1210/jc.2007-2646.
